# Distinct ensembles in the prelimbic cortex track different measures of motivation for cocaine and water reinforcers

**DOI:** 10.64898/2026.05.24.727445

**Authors:** Karla J. Galvan, Sofia D. Grijalva Torres, Rosalie E. Powers, Daniel E. Calvo, Travis M. Moschak

**Affiliations:** Department of Biological Sciences, University of Texas at El Paso, El Paso, TX, 79968, USA

## Abstract

The motivation to pursue drugs is a fundamental element of substance use. Different tasks assess motivation in the face of effort, punishment, or the absence of the drug, and recent studies have suggested that a shared latent variable may drive behavior across these tasks. The prelimbic cortex (PL) is implicated in these behaviors, but it is unknown if its role is driven by shared or distinct neural ensembles. We recorded PL activity using in vivo endoscopic calcium imaging in 32 male and female Sprague-Dawley rats as they completed cocaine or water self-administration, extinction, progressive ratio, and punished self-administration. We found that behavior across tasks was driven by a single latent variable in water, but not cocaine rats. We also found distinct neural populations that tracked reward pursuit. We found that one population of ‘cost-sensitive’ neurons had significantly fewer neurons present during the progressive ratio task. A second population of ‘reward-sensitive’ neurons had significantly fewer neurons present during the extinction task. Individual rats with more of these neurons present during the task had significantly higher levels of reward pursuit in the progressive ratio and extinction tasks, respectively. Furthermore, this relationship was true across cocaine and water rats, suggesting a general role in motivation independent of reward type. When we examined whether shared patterns of neural activity predicted shared patterns of behavior across the tasks, we found no relationships. Thus, our findings suggest that distinct facets of reward motivation are tracked by distinct ensembles in the PL, rather than a shared ensemble.

## 1. Introduction

The motivation to pursue drugs is associated with numerous behavioral traits, such as heightened impulsivity^1–4^, lowered distress tolerance^5,6^, and heightened sign-tracking^7,8^. However, the motivation to pursue drugs is also a fundamental element of substance use disorders in its own right. There are numerous distinct methods that can be used to quantify this. This can include simply assessing the amount of drug consumed, or it can include assessing motivation in the face of effort, punishment, or the absence of the drug itself, among other tests. Furthermore, it has been suggested that these distinct measures may share a single underlying construct that is the primary driver for the motivation to pursue drug^9,10^. However, the existence of such a unifying construct has not been consistently demonstrated. Some studies have demonstrated a shared underlying mechanism for some of the distinct behavioral measures of drug pursuit^11,12^, while others have shown that many of these behaviors are uncorrelated^12–15^. Thus, the degree that these behavioral measures are tapping into a shared underlying motivational construct remains unclear.

One way to assess the presence or absence of an underlying motivational construct is by investigating whether its existence is supported by neural evidence. An appealing target for such an inquiry is the prelimbic cortex (PL). The PL has a well-established role in many aspects of the motivation to seek out drug. It plays a clear role in self-administration^16–19^, extinction^16,17,20^, reinstatement^16,21–23^, effortful pursuit^24,25^ and punished pursuit^26–28^ of drug. Additionally, several studies have demonstrated that different ensembles in the PL differentially encode reward as a function of reward type (e.g. sucrose vs cocaine^17,29,30^). However, it is not known if distinct ensembles in the PL differentially encode these distinct aspects of drug motivation or if there is a shared group of neurons in the PL driving motivation for drug. Establishing whether there is a shared or unique role for the PL in these distinct behavioral measures is a critical step towards developing a general or tailored approach to treatment development.

In the current study, we sought to investigate the role of PL ensembles across multiple measures of drug motivation in rats, with a specific focus on cocaine and water self-administration, extinction, progressive ratio, and punished self-administration.

## 2. Materials and Methods

### 2.1 Animal Housing and Surgery

Thirty-two adult male and female Sprague Dawley rats (60–90 days old; 250–400 g; Envigo) were single-housed in a temperature-controlled room (66°F) with a 12-hour light/dark cycle. Animals were allowed at least one week of acclimation before handling and had ad libitum access to food (Purina Laboratory Chow) and water. All experiments were conducted in accordance with the National Institutes of Health *Guidelines for the Care and Use of Laboratory Animals* and approved by The University of Texas at El Paso.

After one to two weeks of acclimation, rats underwent surgery for unilateral viral infusion (GCaMP6s) and lens implantation into the prelimbic cortex (coordinates: AP +2.7, ML ±0.6, DV –3.6 for viral infusion and –3.4 for lens implantation), alternating left and right hemispheres from bregma.

Anesthesia was induced using a mixture of ketamine hydrochloride (100 mg/kg) and xylazine (10 mg/kg), with supplemental isoflurane (Covetrus, ANADA 200-237) administered as needed to maintain anesthesia. Following surgery, animals were allowed a 5-day recovery period.

Six weeks post-surgery, GCaMP expression was verified using the Miniscope software system (UCLA Miniscope, Los Angeles, CA). Upon confirmation of cellular fluorescence, rats were implanted with miniscope baseplates to secure the cameras used for recording neuronal activity during behavior. Subsequently, each animal was implanted with a chronic indwelling catheter (Access Technologies, Skokie, IL, USA) in the right jugular vein. The catheterization procedure followed previously described methods^31,32^.

### 2.2 Apparatus

Cocaine or cocaine-histamine infusion and water delivery were controlled by a syringe pump (Med Associates). For infusions, tubing was tethered using a counterweighted arm to allow free movement and connected to the back-mounted catheter. For water administration, a syringe pump was connected to a water dish. The chamber (10 × 10 × 12 in) contained two nose-poke ports and a cue light positioned above the apparatus.

Each time a rat self-administered cocaine or water, a tone was activated, the entire chamber was illuminated, and the solution was dispensed. All behavioral data were collected using MedPC software.

#### 2.2.1 Endoscopic In Vivo Calcium Imaging

Imaging was performed using the Miniscope system (UCLA Miniscope, Los Angeles, CA) synchronized with task events. The Miniscope was attached to the implanted baseplate and connected via a tether to a commutator, preventing tangling and allowing free movement. The opposite end of the tether was connected to a DAQ (Data Acquisition device), which received TTL (Transistor-Transistor Logic) signals from the MED-PC system and was linked to a laptop. Neuronal activity was visualized and recorded in real time using Miniscope software. Recordings were segmented into alternating intervals: 10 minutes with the LED on, followed by 5 minutes with the LED off, to minimize photobleaching. Calcium imaging was performed during all relevant behavioral sessions.

Miniscope data were processed as we have done previously^33^. Raw video files were downsampled and subsequently processed using CaImAn^34,35^. Following motion correction, potential neurons were identified based on the spatial/temporal signature of each calcium signal. All such neurons were inspected for artifactual or noise contamination and only retained if they passed visual threshold. Finally, CaImAn was used to coregister neurons across tasks. MATLAB and Neuroexplorer were used to align individual traces to relevant events in the tasks.

### 2.3 Behavioral Procedures

#### 2.3.1 Self-Administration

Before neural recording, animals completed 2 weeks of self-administration followed by one month of abstinence, as well as several food-based operant tasks in contextually distinct operant chambers using different operands (lever press vs. the nosepokes used in the current study) that were part of a different study. Other than the self-administration task, these tasks were distinct from the current study and are not reported on here. During the self-administration task, rats inserted their noses into an illuminated (cue) port to either receive 0.6 ml/kg water from a dispenser or an intravenous dose of cocaine (1 mg/kg/infusion). Each reward delivery was accompanied by illumination of the chamber and a tone. A 20-second timeout period followed each administration, during which no additional infusions could be obtained. Rats were water-restricted: males in the water group received 20 mL per day, females 15 mL; males in the cocaine group received 30 mL, and females 25 mL. During the neural recording, the duration of the self-administration task was 2 hr. Number of reinforcers earned was the dependent measure in this task.

#### 2.3.2 Extinction

In this task, drug-paired cues (tone and light) were presented, but neither cocaine nor water was delivered. Sessions lasted 2 hours. Number of nosepokes made was the dependent measure in this task.

#### 2.3.3 Progressive Ratio

Following extinction, rats were returned to the self-administration chamber the next day for a 2-hour session to re-establish the association between cues and reward delivery. The following day, rats were again placed in the chamber to perform a progressive ratio task according to the formula 5e^0.2n^-5 describing the number of responses required on trial n. Final completed ratio was the dependent measure in this task.

#### 2.3.4 Punished Self-Administration

After the progressive ratio task, rats were again returned to the operant chamber for a 2-hour re-acquisition session. The next day, they were tested in a fixed-ratio task for self-administration of a cocaine-histamine mixture (1 mg/kg/infusion cocaine, 5 mg/kg/infusion histamine). Histamine causes an acute delocalized itching sensation and suppresses cocaine intake^36^. Number of reinforcers earned was the dependent measure in this task. Finally, it is important to note that only cocaine rats were tested in this task, and that water rats were not tested in this task.

### 2.4 Data Analysis

#### Behavior

Our dependent measures were Reinforcers (Self-administration and Punished Self-administration), Nosepokes (Extinction), and Completed Ratio (Progressive Ratio). For each task, we used a 2 x 2 ANOVA with Drug and Sex as a factor to determine differences in behavioral responding (Punished Self-administration only had a t-test since there were no water rats tested in that task). We also ran two Principal Component Analyses on Water and Cocaine rat data across all tasks to determine if underlying latent variables described the data.

#### Neural Activity and Classification

In prior studies we statistically classified neural activity according to the change in of activity at a given event^37,38^. That method was not feasible in the current study as some animals had very few trials in which they responded, which would have biased phasic neurons towards those rats which had a high response rate. Therefore, to categorize neurons into different patterns of activity, we performed Uniform Manifold Approximation and Projection (UMAP) Analysis on perievent histograms aligned around reinforcer nosepokes across all tasks. As clear clusters did not emerge, we classified neurons into 4 categories (Type 1, 2, 3, and 4) according to the 10% most extreme edges of the two emergent UMAPs. We then analyzed perievent histograms for all neurons grouped in one of the four categories. We also assessed the proportion of neurons that fell into each category for a given rat.

#### Relative proportion of neural types across tasks

To determine the relative proportion of neurons across tasks and account for the unbalanced nature of the dataset (some animals had missing neural activity in one or more tasks), we ran linear mixed-effects models. Repeated observations across tasks per rat were nested within-subject, and subject was included as a random effect to account for this within-subject dependence. We also incorporated between-subject information for those rats that only completed a subset of tasks. Our factors were Drug (Water or Cocaine), Sex (Female or Male), and Task (Self-Administration, Extinction, and Progressive Ratio). To accommodate the fact that our Punished Self-Administration task only had rats in the cocaine group, we also ran the same analysis on only cocaine rats but with Punished Self-Administration as an additional level of Task. Subsequently, we ran Pearson correlations between neural activity and behavior.

We also used PCA on neural data (relative proportion of neural types across tasks) to investigate if there were latent variables driving shared activity. We only analyzed extinction and progressive ratio data (8 variables comprising each of the 4 neuron types for the 2 tasks), because including self-administration and punished self-administration data would have reduced the sample size to a point where PCA would not be feasible.

#### Shared links between neural activity and behavior

To assess the shared links between neural activity and behavior, we directly predicted latent behavior variables from latent neural variables. Because we were only able to input extinction and progressive ratio data into the neural analysis, we ran a second PCA on behavioral data that only included extinction and progress ratio data. We then ran either multiple regression (on those data sets that only had one dependent variable) or canonical correlation analysis (on those data sets with two dependent variables) with neural latent variables predicting behavioral latent variables.

#### Shared neural identity analysis

We also sought to determine if neural categories persisted across different tasks. Using neurons coregistered across tasks, we assessed the tendency for neurons to retain the same neural type across tasks. To do this, for each rat we calculated the percent of neurons in one task that remained in the same category in another task (e.g. is a Type 1 neuron in Self-admin a Type 1 neuron in Extinction), as well as the percent of neurons that switched categories. We then subtracted these two percentages to obtain a preference score for retain the same or different classification. Finally, we ran Pearson correlations between preference scores across different pairs of tasks.

## 3. Results

### Behavior

We first assessed differences in responding as a function of sex and drug for each of the tasks. Females in the water group obtained significantly more reinforcers than other rats during the self-administration task (Sex x Drug interaction: F(1,25) = 10.44, p = 0.003; Fig. 1A). Females had significantly higher nosepokes under extinction across groups (main effect of Sex: F(1,30) = 6.27, p = 0.018; Fig. 1B) and there was a trend for higher nosepokes under extinction in the cocaine group (main effect of Drug: F(1,30) = 3.99, p = 0.055; Fig. 1B). Finally, rats had significantly higher breakpoints for cocaine than for water (main effect of Drug: : F(1,25) = 20.83, p < 0.001; Fig. 1C). There were no other significant relationships between sex and drug.

**Figure 1.**
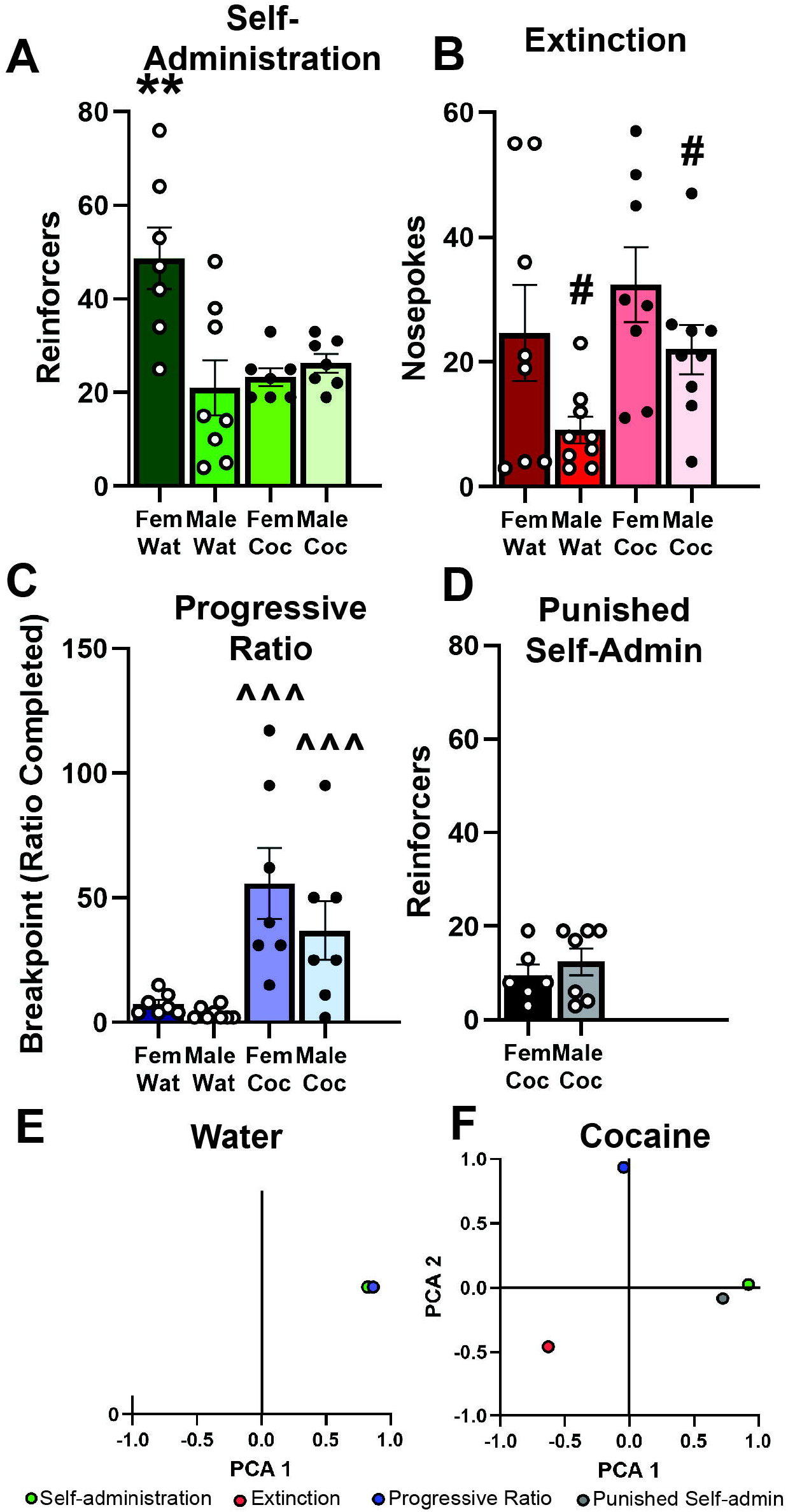
Behavior. **A)** Female water rats earned significantly more reinforcers that other groups. **B)** Female rats had significantly more responses that males under extinction. **C)** Cocaine rats had significantly higher breakpoints than water rats under progressive ratio. **D)** There were no differences between sex in the number of reinforcers earned under punishment (no water rats were run in this task). **E)** In water rats, behavior across tasks tightly loaded onto one principal component accounting for 83.9% of the variance. **F)** In cocaine rats, behavior was less related, with only self-administration and punished self-administration showing any relationship. ** p < 0.01 comparing female water rats to others, # p < 0.05 comparing male to female rats, ^^^ p < 0.001 comparing cocaine to water rats.

To determine if a shared latent variable drove motivation for either water or cocaine across the tasks, we ran principal component analyses (PCAs) separately on water and cocaine rats. For water rats, responding across all three tasks was strongly driven by one underlying variable which accounted for 83.9% of the variance (Fig. 1E). Conversely, responding for cocaine split into two variables accounting for 45.9% and 25.8% of the variance. Responding under self-administration and punished self-administration were linked, but the other two behaviors were unrelated (Fig. 1F). Thus, there appeared to be a shared latent variable driving motivation for water, but not cocaine.

### Prelimbic activity

To categorize neurons into different patterns of activity, we performed UMAP Analysis on perievent histograms aligned around reinforcer nosepokes across all tasks. As clear clusters did not emerge, we classified neurons into 4 categories (Type 1, 2, 3, and 4) according to the 10% most extreme edges of the two emergent UMAPs (Fig. 2A) which revealed distinct firing patterns (Fig. 2B). There were no significant differences in firing patterns for each of the neural categories across each of the behaviors (Fig. 2C-J).

**Figure 2.**
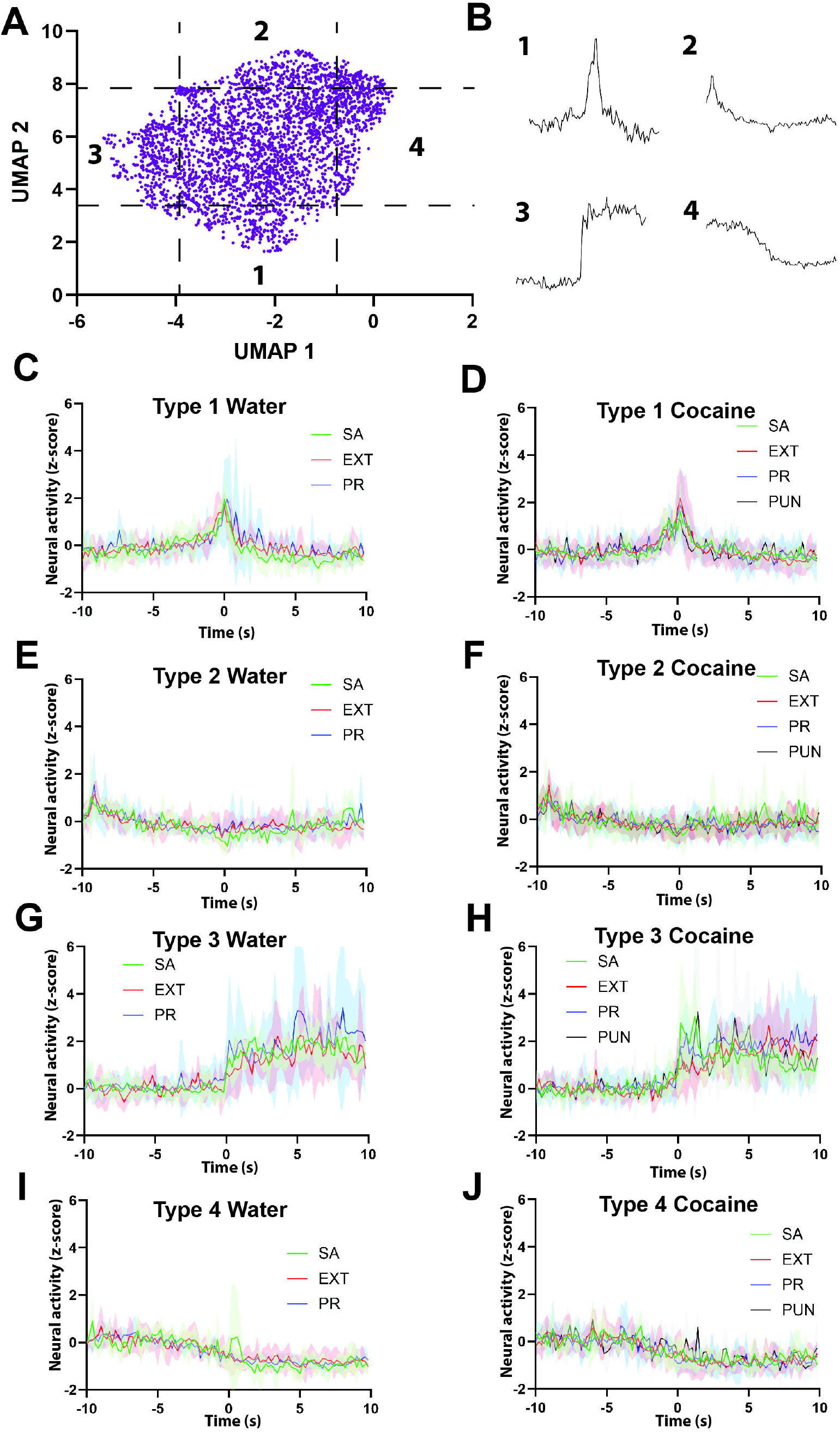
Neural classification. **A)** UMAP data for each neuron. As there were no clear clusters, we took the 10% most extreme data points from the two UMAPs and categorized them as neuron Types 1-4. **B)** The average perievent histogram for neuron Types 1-4. We saw no differences in the perievent histograms for each neuron type across task and drug (**C-J**).

In addition to examining differences in firing patterns, we also examined the relative proportion of neurons for each category present in each of the behavioral tasks. There were significantly fewer Type 1 neurons in the progressive ratio task, and there trended to be fewer in the punished self-administration task (Main effect of task (including water rats but excluding punished self-admin): F(2,33.23) = 9.86, p < 0.001; Main effect of task (including punished self-admin but excluding water rats): F(3,461.0) = 12.12, p < 0.001; Fig. 3A). Furthermore, there were significantly fewer Type 3 neurons in the extinction task (Main effect of task (including water rats but excluding punished self-admin): F(2,11.73) = 11.18, p = 0.002; Main effect of task (including punished self-admin but excluding water rats): F(3,5.21) = 8.67, p = 0.018; Fig. 3C). Additionally, female water rats had significantly fewer Type 3 neurons during the progressive ratio than other rats (Task x Sex x Drug interaction (including water rats but excluding punished self-admin): F(2,11.73) = 5.46, p = 0.021). There were no differences in Type 2 or Type 4 neurons (Fig. 3B,D). Other than the one aforementioned interaction with female water rats, there were no differential effects of drug or water on the percent on neurons in each category across tasks. This suggests that the role these neurons play may be generalizable across certain rewards (here, water and cocaine).

**Figure 3.**
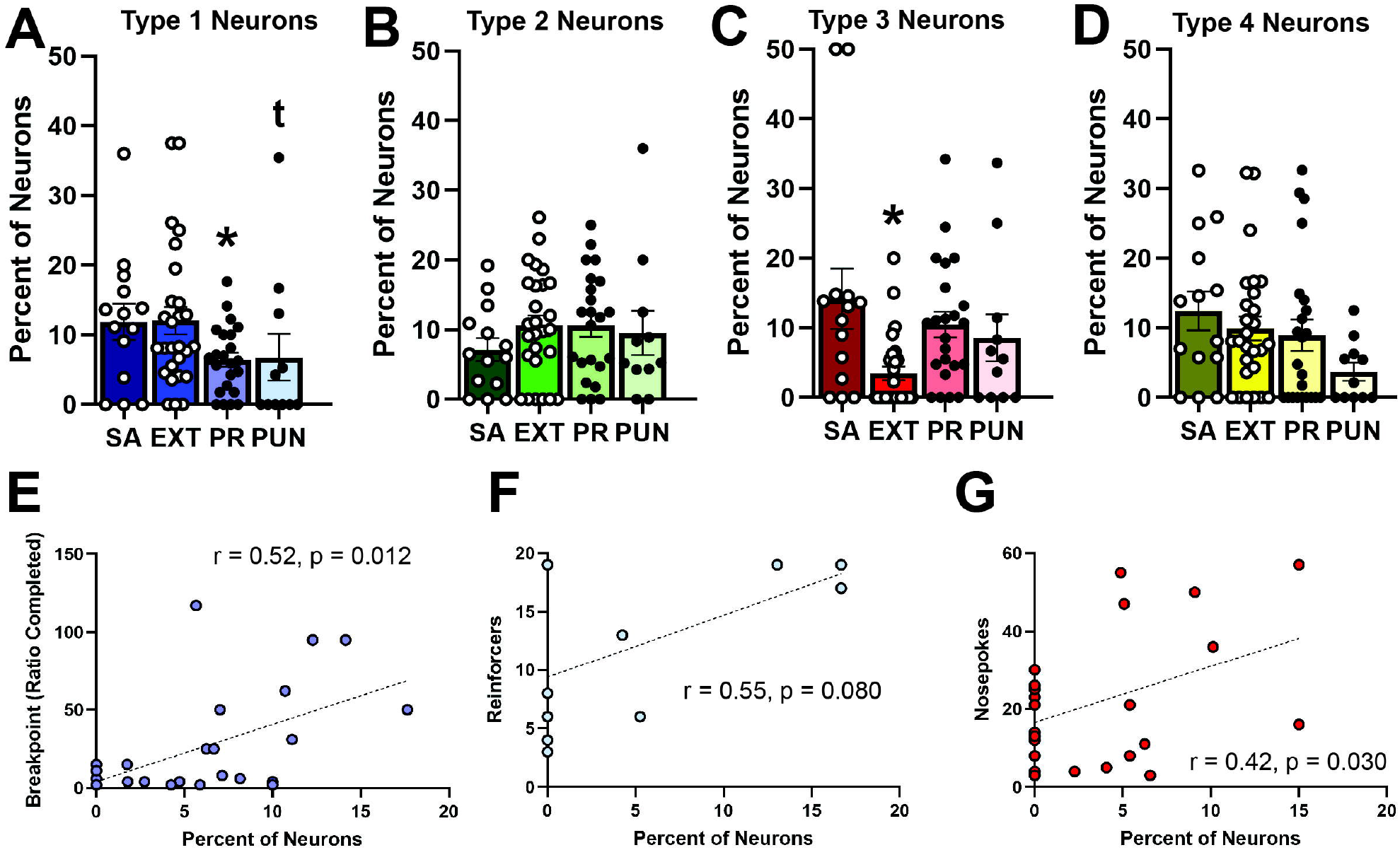
Prevalence of neural type across task. **A)** There were significantly fewer Type 1 neurons in the progressive ratio task and a trend for fewer in the punished self-admin task. **B)** There were no differences in the relative prevalence of Type 2 neurons across tasks. **C)** There were significantly fewer Type 3 neurons in the extinction task. **D)** There were no differences in the relative prevalence of Type 4 neurons across tasks. **E)** Rats with more Type 1 neurons during the progressive ratio task had significantly higher breakpoints. **F)** Rats with more Type 1 neurons during the punished self-administration task tended to earn more reinforcers. **G)** Rats with more Type 3 neurons during the extinction tasks had significantly higher nosepokes. * p < 0.05, t p < 0.10

Since the presence of Type 1 neurons dropped during costs (either effort or punishment) and the presence of Type 3 neurons dropped in the absence of reward (extinction), we label these neurons as “cost-sensitive” and “reward-sensitive”, respectively. To establish how cost-sensitive neurons tracked motivation for reward, we correlated percent of neurons with breakpoint or reinforcers in the progressive ratio and punished self-admin tasks. The percent of cost-sensitive neurons significantly predicted breakpoint (r = 0.52, p = 0.012; Fig. 3E), and trended to predict number of reinforcers obtained under punishment (r = 0.55, p = 0.080; Fig. 3F). We performed a similar analysis for reward-sensitive neurons and extinction, and found that the percent of reward-sensitive neurons significantly predicted nosepokes under extinction (r = 0.42, p = 0.030; Fig. 3G). Thus, the relative absence or presence of cost-sensitive and reward-sensitive neurons influences motivation for reward in the domain-relevant tasks.

We also ran PCA on prelimbic data from cocaine and water rats to assess if there was a shared latent variable across tasks. We focused on the percent of neurons that classified as each type (i.e. the data comprising Fig. 3) as this was the most behaviorally relevant phenotype. We also only analyzed extinction and progressive ratio data (8 variables comprising each of the 4 neuron types for the 2 tasks), since including self-administration and punished self-administration data would have reduced the sample size to a point where PCA would not be feasible. We found that for both water and cocaine rats, three latent variables captured the majority of the data (water: 77.71%, cocaine: 69.44%). Because there were no differences between water and cocaine rats in neural activity, we also ran the PCA combined across the two groups and found that three latent variables again captured the majority of the data (57.79%).

#### Shared neural activity

Finally, we sought to determine if shared neural activity across tasks predicted the shared aspects of behavior that we saw in Fig. 1E,F. Because we were only able to input extinction and progressive ratio data into the neural analysis, we ran a second PCA on behavioral data that only included extinction and progress ratio data. We found that PCA of both water and cocaine behavioral data yielded a single latent variable (water: 78.22%; cocaine: 69.01%); PCA of water and cocaine combined did not yield any latent variable that explained more variance than the variables themselves. Therefore, we analyzed a multiple regression for the two separate analyses and a canonical correlation analysis for the water + cocaine analysis. For all three analyses, the shared neural activity did not predict the shared behavioral activity (water: adjusted R^2^ = 0.22, p = 0.214; cocaine: adjusted R^2^ = -0.21, p = 0.744; water + cocaine: canonical correlation r = 0.61, p = 0.293; Fig. 4A-C). Thus, prelimbic activity appears to have distinct roles across behaviors (Fig. 3) rather than a shared role (Fig. 4).

**Figure 4.**
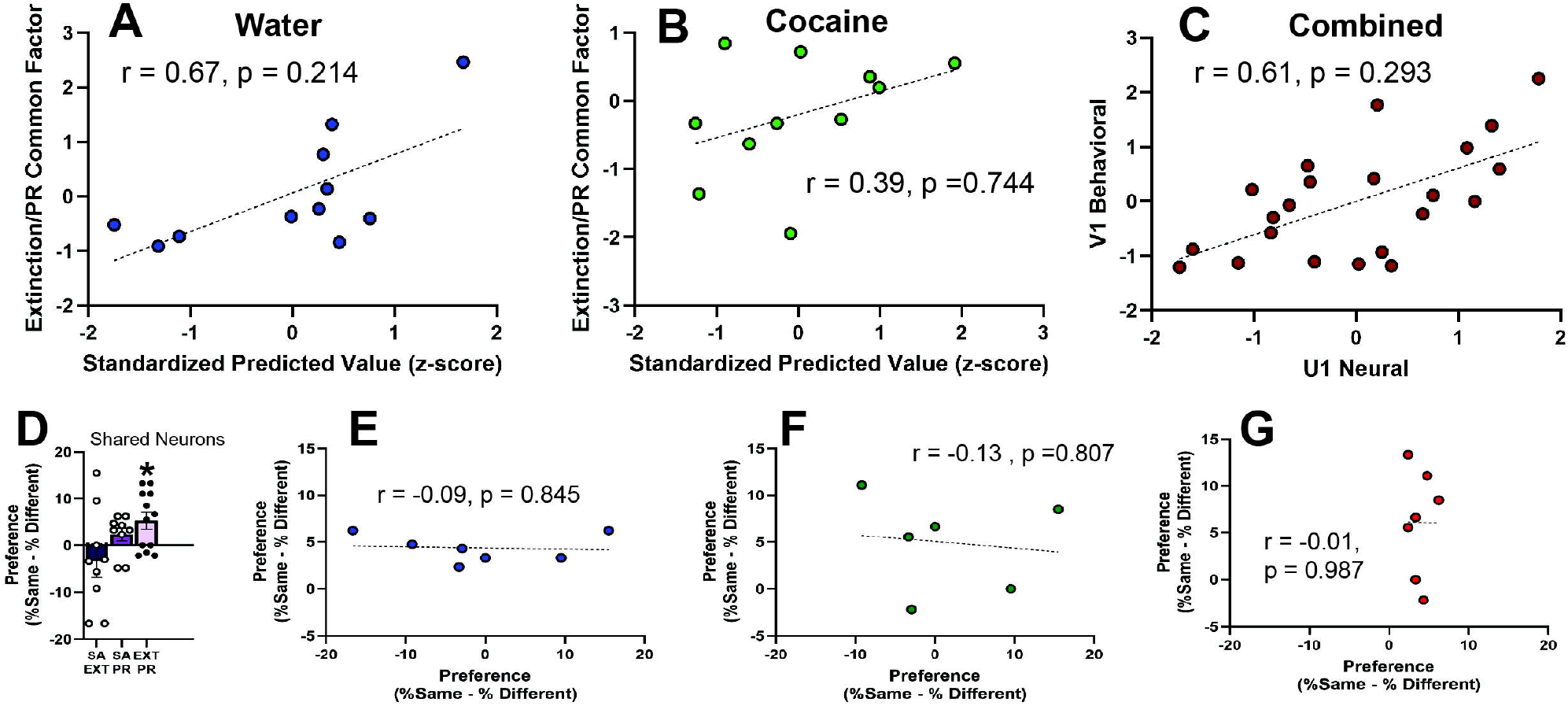
Shared activity. Shared neural activity did not predict shared behavioral activity for either water (**A**), cocaine (**B**), or a combination of the two (**C**). **D)** Neurons classified as a given category in the extinction task were significantly more likely than chance to be classified as the same category in the progressive ratio task. This was not true for other task pairings. **E-G)** The tendency for individual rats to share neural classifications across a given two tasks did not predict the same tendency in other tasks. * p < 0.05

We also sought to determine if neural categories persisted across different tasks. Using neurons coregistered across tasks, we assessed the tendency for neurons to retain the same neural type across tasks. We found that extinction and progressive ratio tasks had a significant tendency to share neural activity across tasks (t(11) = 2.97, p = 0.013; Fig. 4D). Based on prior unpublished work, we also assessed whether a tendency the share neural activity across a given set of tasks would predict a tendency to share neural activity across a different set of tasks, but we found no significant relationships (Fig. 4E-G). Thus, while there is some shared activity across tasks, this is unique to the given tasks and does not predict behavior in those tasks.

### Histology

All placements were verified with a rat brain atlas^39^ (Fig. 5). Rats with placements outside the prelimbic cortex were excluded from the analysis.

**Figure 5.**
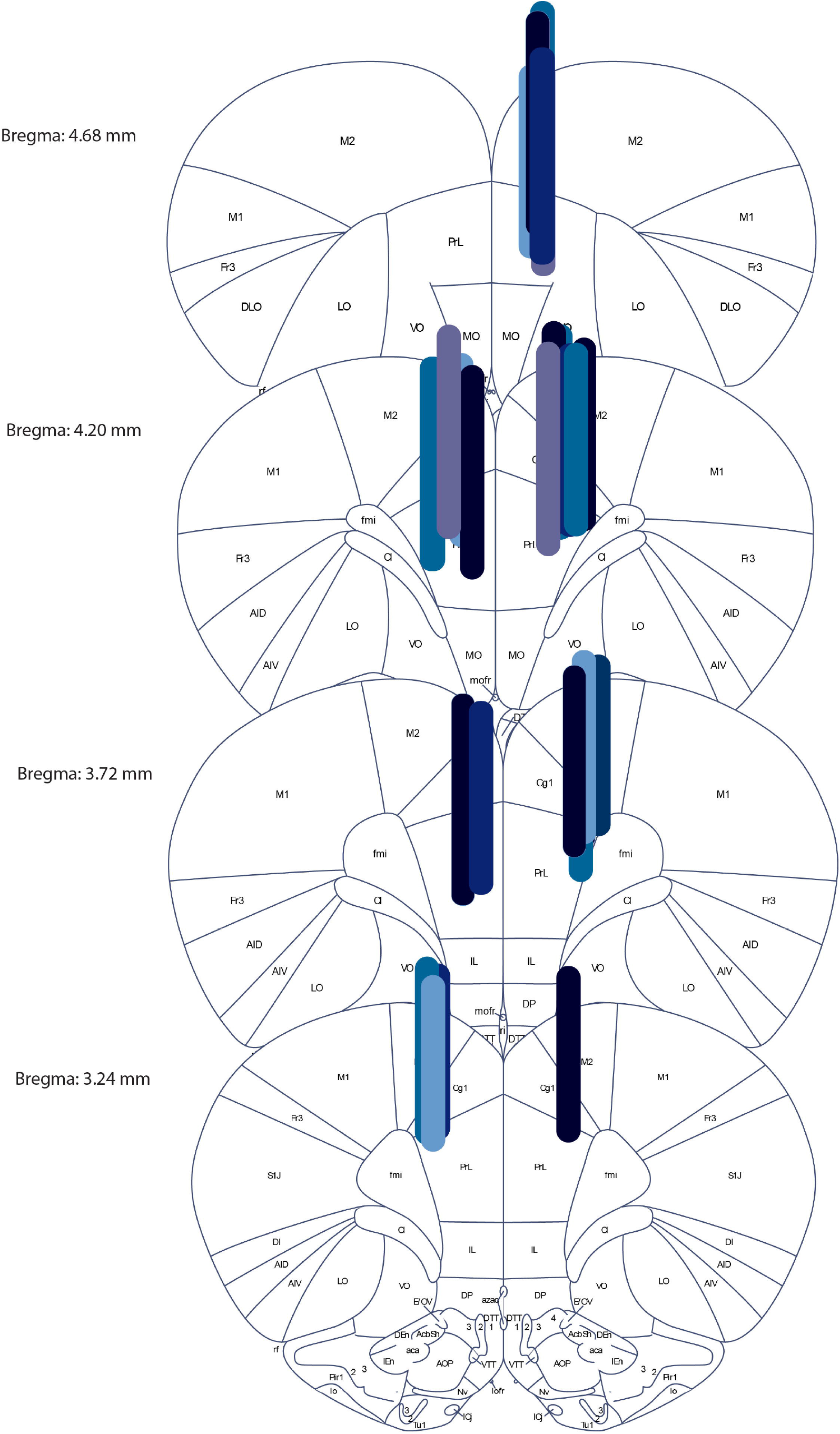
Location of placements in the prelimbic cortex.

## 4. Discussion

Several studies have investigated the relationships between different measures of drug motivation, but few have investigated how neural activity tracks those relationships. Here, we investigated the role of PL activity across four distinct tasks of motivation for either cocaine or water: self-administration, extinction, progressive ratio, and punished self-administration. Behaviorally, we found that performance on these tasks was highly correlated in water rats, but not in cocaine rats. Analysis of neural activity established distinct populations of neurons that differentially tracked behavior. Notably, the relative number of neurons in some of these distinct populations was lower in some tasks, and the degree to which is was lowered predicted reward motivation. Thus having established unique contributions of different neural populations to behavior, we also assessed if there were shared neural predictors of behavior, but found none. Therefore, our data suggest that PL activity tracks unique, but not shared aspects of reward motivation.

Our behavioral findings with cocaine have some support in existing literature. We found that self-administration and punished self-administration loaded similarly onto the same latent variable, while this was not seen for extinction or progressive ratio. The link we found between cocaine self-administration and punished cocaine self-administration has been previously reported^11,12^, and cocaine extinction has explicitly been established as loading onto a separate factor^12,15^. However, unlike our study, other studies have demonstrated that progressive ratio for cocaine was linked to punished cocaine self-administration^11,12^, although at least one other study agrees with our findings in suggesting that the two are separable^13^. Thus, our findings reinforce prior studies suggesting that extinction tracks a distinct process, while adding to the mixed body of evidence on effortful and punished cocaine motivation. Our findings with water were unexpected and novel, as we found that water self-administration, extinction, and progressive ratio were all strongly driven by the same underlying latent variable. The distinct measures of motivation for natural reward have primarily been viewed as separate processes^9,10^, although to our knowledge no study has explicitly assessed the relationships between them and little work has investigated the use of water as a reward. Our data suggests that, for water, these measures are tapping into a shared motivational construct. Thus, in our hands, the motivation to pursue either cocaine or water follows very distinct patterns.

Our neural findings support and expand upon existing literature demonstrating a role for the PL in motivation for drug and natural reward^16–30^. Notably, we found distinct neural populations in the PL that were differentially suppressed as a function of the task used to assess behavior. Specifically, the presence of “cost-sensitive” neurons was lower in the progressive ratio task and tended to be lower in the punished self-administration task, while the presence of “reward-sensitive” neurons was lower in the extinction task. Furthermore, the relative prevalence of these neurons predicted behavioral performance in the relevant tasks: more “cost-sensitive” neurons predicted greater breakpoint in the progressive ratio task, and more “reward-sensitive” neurons predicted more nosepokes under extinction. These findings occurred in both cocaine and water animals, suggesting that this was a general feature of reward. In total, these results suggest that independent PL ensembles track different measures of motivation for reward. The independent nature of these ensembles was further confirmed when we found no shared relationships between neural activity and behavior across the tasks.

These findings expand upon the relatively sparse literature examining neural ensembles across measures of motivation. Prior studies have demonstrated the existence of separate PL ensembles for self-administration and extinction^15,40^, although another study found significant PL overlap across the two behaviors^17^. Regardless, ours is the first to extend this to progressive ratio and punished self-administration. Furthermore, Although it is likely that cocaine and water recruited distinct PL ensembles in these tasks (as suggested by^29,41,42^), it is notable that there were no differences between the two groups in their recruitment of the cost-sensitive and reward-sensitive neurons across tasks. Thus, while these tasks appear to recruit distinct ensembles, the nature of that recruitment may occur independently of the reward type.

In total, our study suggests that distinct ensembles in the PL track distinct measures of motivation for reward, independent of whether that reward is cocaine or water. Future studies should establish the causal independence of these ensembles, and expand into other cortical and subcortical regions that have been implicated in motivation for reward.

## Acknowledgements

This work was supported by and National Institute of Health (NIH) grant U54MD007592-28, National Institute on Drug Abuse (NIDA) grant DA045764, and NIDA grant DA058653 awarded to T.M.M. S.D.G.T. was supported by National Institute of General Medical Sciences (NIGMS) grant GM145529. The authors would also like to thank Serina Batson and Jaylene Scales for technical assistance. Imaging was performed at the Imaging & Behavioral Neuroscience (IBN) Facility Imaging Core at The University of Texas at El Paso, supported by the College of Science and the Office of Research & Innovation. The IBN Facility was supported by the Office of the Director, National Institutes of Health (NIH), under award number C06OD032074. We thank Sivasai Balivada for his assistance. The content is solely the responsibility of the authors and does not necessarily represent the official views of the NIH.

